# A high-fat/high-protein, Atkins-type diet exacerbates *Clostridioides (Clostridium) difficile* infection in mice, whereas a high-carbohydrate diet protects

**DOI:** 10.1101/834903

**Authors:** Chrisabelle C. Mefferd, Shrikant S. Bhute, Jacqueline R. Phan, Jacob V. Villarama, Dung M. Do, Stephanie Alarcia, Ernesto Abel-Santos, Brian P. Hedlund

## Abstract

*Clostridioides difficile* (formerly *Clostridium difficile*) infection (CDI) can result from the disruption of the resident gut microbiota. Western diets and popular weight-loss diets drive large changes in the gut microbiome; however, the literature is conflicted with regard to the effect of diet on CDI. Using the hypervirulent strain *C. difficile* R20291 (RT027) in a mouse model of antibiotic-induced CDI, we assessed disease outcome and microbial community dynamics in mice fed two high-fat diets in comparison with a high-carbohydrate diet and a standard rodent diet. The two high-fat diets exacerbated CDI, with a high-fat/high-protein, Atkins-like diet leading to severe CDI and 100% mortality, and a high-fat/low-protein, medium-chain triglyceride (MCT)-like diet inducing highly variable CDI outcomes. In contrast, mice fed a high-carbohydrate diet were protected from CDI, despite high refined carbohydrate and low fiber content. 28 members of the *Lachnospiraceae* and *Ruminococcaceae* decreased in abundance due to diet and/or antibiotic treatment; these organisms may compete with *C. difficile* for amino acids and protect healthy animals from CDI in the absence of antibiotics. Together, these data suggest that antibiotic treatment might lead to loss of *C. difficile* competitors and create a favorable environment for *C. difficile* proliferation and virulence that is intensified by high-fat/high-protein diets; in contrast, high-carbohydrate diets might be protective regardless of the source of carbohydrate.

## Introduction

*Clostridioides difficile* (formerly *Clostridium difficile*) is an endospore-forming member of the phylum *Firmicutes* that is the leading cause of antibiotic-associated and hospital-acquired diarrhea. *C. difficile* infections (CDIs) make up > 70% of healthcare-associated gastrointestinal infections, with symptoms ranging from mild diarrhea to severe infections resulting in ulcerative colitis and toxic megacolon (1). Moreover, CDI is financially taxing on U.S. hospital management (2) and is the cause of over 500,000 infections and 29,000 deaths annually, according to a 2015 report (3).

Stable and complex microbial communities in the gut act as a natural barrier against *C. difficile* (4), but broad-spectrum antibiotics can disrupt the native microflora, allowing *C. difficile* to multiply and cause CDI (5). Importantly, *C. difficile* has innate resistance to multiple antibiotics and CDI is closely linked to administration of ampicillin, amoxicillin, cephalosporins, clindamycin, and fluoroquinolones (6). In order to cause successful infection, *C. difficile* spores must germinate, grow within the intestinal lumen, and produce toxins that mediate tissue damage and inflammation (7). Specific chemical signals are needed for each of these steps. For example, spore germination is promoted by variety of amino acids and primary bile salts, but inhibited by secondary bile salts (8); growth can be supported by fermentation of amino acids or carbohydrates, but is inhibited by short-chain fatty acid (SCFA), products of carbohydrate fermentation (9–11); finally, toxin production is inhibited by several amino acids, particularly cysteine (12). Thus, it is logical that diet might affect the incidence and severity of CDI, yet the literature is contradictory on relationships between diet and CDI.

Several studies have suggested that high-carbohydrate/low protein diets can mitigate antibiotic-induced CDI. Moore *et al.* (13) hypothesized that poor nutrient status would worsen CDI, but instead found that protein-deficient diets (with increased carbohydrate content) mitigated CDI severity in C57BL/6 mice infected with the hypervirulent strain VPI10463 (Ribotype 078 (RT078)). Another study using humanized mice inoculated from antibiotic-induced dysbiotic subjects reported increased lumen amino acid concentrations and severe CDI compared with those inoculated from control subjects (11). The same study showed that *C. difficile* strain 630 expressed the proline reductase, PrdA, only in dysbiotic mice and that *prdB* mutants unable to use proline as the Stickland electron acceptor failed to colonize mice. Finally, they showed that low-protein and specifically low-proline diets lessened colonization and virulence. A separate study found that mixtures of microbiota-accessible carbohydrates (MACs), or specifically inulin, decreased *C. difficile* burdens in humanized mice, while stimulating growth of carbohydrate-utilizing microbes and SCFA production (10).

In contrast, other studies have implicated carbohydrates, specifically simple sugars, in the proliferation of hypervirulent, epidemic *C. difficile* strains. One study reported on the independent evolution of mechanisms to utilize the artificial sweetener trehalose in RT027 and RT078 and showed that trehalose supports growth of these ribotypes *in vitro* (14). Trehalose also increased toxin production and decreased survival in a humanized mouse model of CDI but did not increase *C. difficile* burden. Similarly, Kumar and colleagues reported positive evolutionary selection on the fructose phosphotransferase system and several other enzymes involved in the transport and fermentation of simple sugars in recently evolved strains of *C. difficile*, including RT027 (15). That study also reported that glucose and fructose enhanced growth and sporulation of RT027 *in vitro* and shedding in a mouse model of CDI, but relationships between dietary monosaccharides and virulence were not reported.

Western diets would seem to favor CDI since they are enriched in both protein and simple sugars yet deficient in MACs and other fiber sources (16), and indeed rates of CDIs are highest in developed countries (17). Modern weight-loss diets such as Atkins and ketogenic diets are extreme because the majority of calories are from fat and protein, and carbohydrates typically contribute less than 10% of caloric intake (18, 19). These diets have been wildly popular; for example, Dr. Atkins’ *Diet Revolution* is the best-selling diet book in history (20). Keto diets are similar to Atkins’ diets, but tend to be more extreme, reducing both dietary carbohydrates and proteins. Yet, despite the increasing evidence tying *C. difficile* evolution and pathogenesis to diet, and the continuing revolution of modern diets with extreme macronutrient composition, diet is not featured as a major factor in models of CDI (17).

Here we assessed the effect of diet, including a high-fat/high-protein Atkins-like diet, a high-fat/low-protein keto-like diet resembling the medium-chain triglyceride (MCT) diet, and a high-carbohydrate diet, on the outcome of antibiotic-associated CDI using the hypervirulent *C. difficile* R20291 and described concomitant changes in microbial community diversity and composition.

## Results

### A high-fat/high-protein diet exacerbates CDI, yet a high-carbohydrate diet protects

To determine whether diet affects the progression of CDI, groups of five mice were fed diets varying in macronutrient composition (Fig. 1): a high-fat/high protein diet, a high-fat/low-protein diet, a high-carbohydrate diet, and a standard lab diet (Files S1–3 and Table S1). To quantify the effect of the diets on CDI severity, morbidity and mortality were examined over the course of the experiment (Fig. 2) using established metrics (21) with amendments, as described in Methods. All infected mice fed the standard lab diet developed moderate CDI signs but eventually recovered. In contrast, only two of the mice fed the high-carbohydrate diet exhibited mild symptoms, and they quickly recovered. The rest of the animals in this group never developed any CDI signs. Mice fed the high-fat/low-protein diet showed CDI symptom onset heterogeneity. Two animals developed severe CDI and became moribund. Meanwhile, three animals developed moderate CDI signs similar to the standard diet and recovered within a week post-infection. Strikingly, all mice fed the high-fat/high-protein diet developed severe CDI signs and were euthanized within four days following *C. difficile* challenge. The difference in survival between mice fed the high-fat/high-protein diet and all other animal groups was significant (*p* = 0.003, log-rank test). Surviving animals in all groups resolved all CDI signs (score of 0) within eight days and were remained healthy for the remainder of the experiment. A control cohort of uninfected mice fed the standard lab diet remained healthy and showed no CDI symptoms for the duration of the experiment (not shown).

**Fig. 1.**
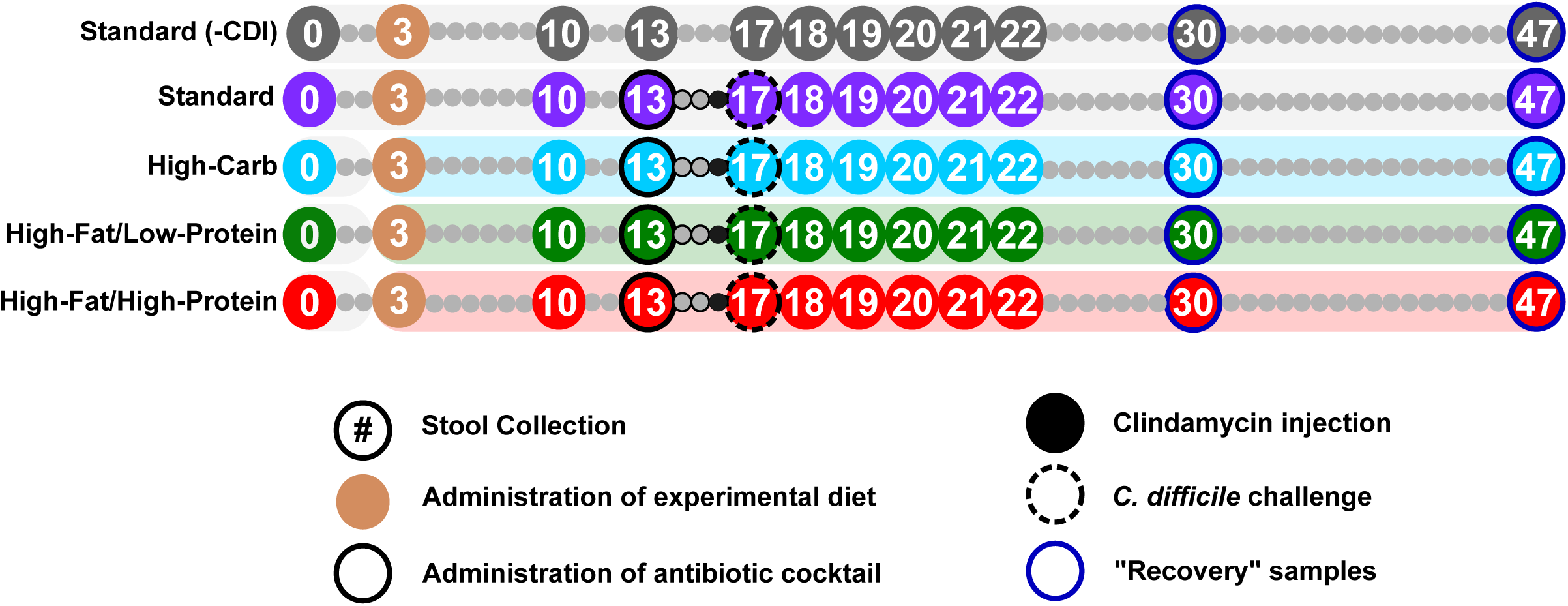
Experimental timeline and stool collection. High-carbohydrate (blue), high-fat/low-protein (green), or high-fat/high-protein (red) diets were introduced on Day 3. An antibiotic cocktail (solid outline) and clindamycin (black-filled circles) were given on Day 13 and Day 16, respectively. Mice were challenged with *C. difficile* R2027 spores on Day 17 (dashed outline). Circles with numbers indicate days fecal samples were collected. Stool collection took place prior to manipulation of mice or experimental treatment.

**Fig. 2.**
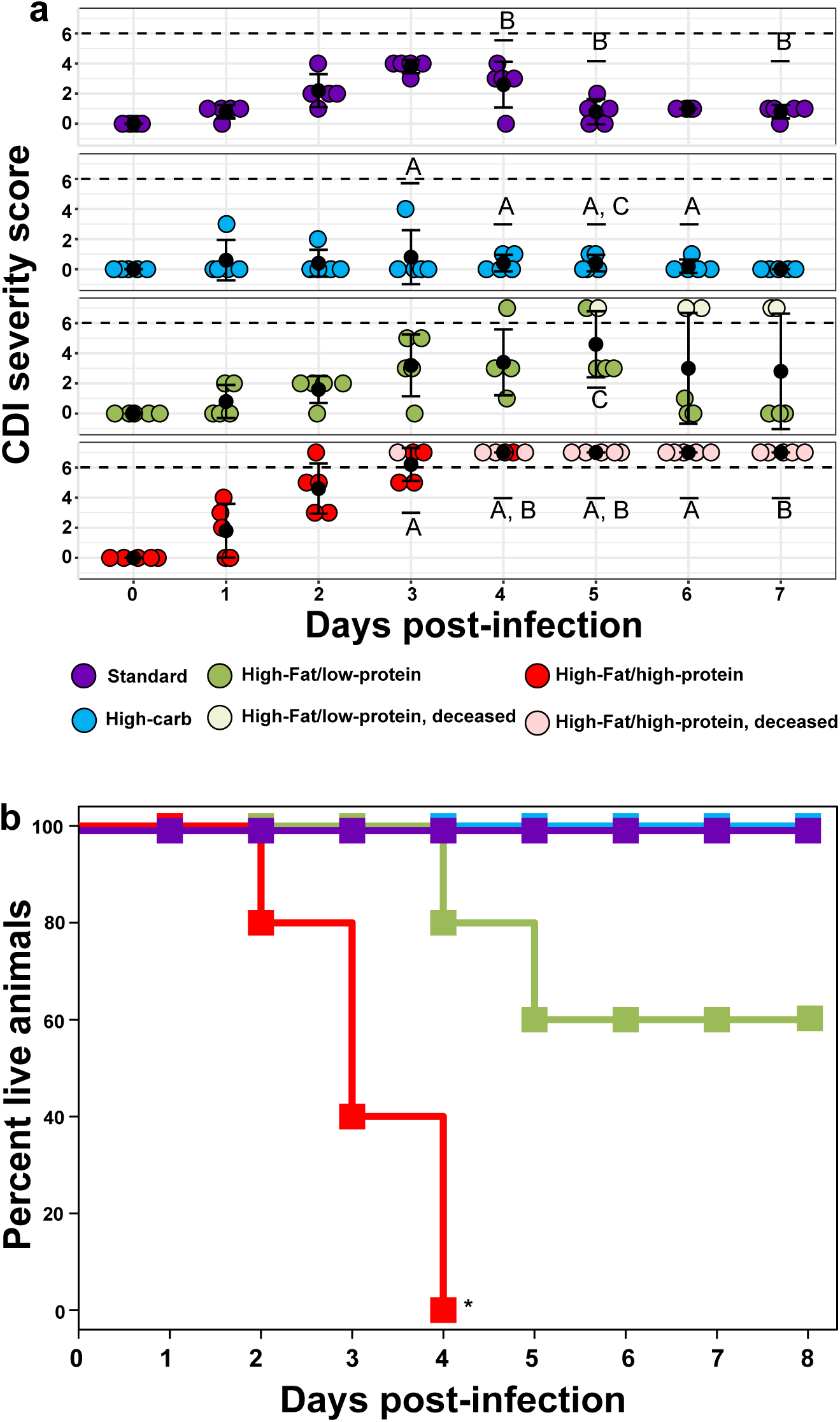
Effect of diet on mice survival and CDI severity post-infection. a) Mean disease severity scores (black dots) for each diet following CDI challenge, with 25^th^ and 75^th^ percentiles shown. Colored dots represent severity scores for individual mice. Dashed lines represent a score of 6, the clinical endpoint. Groups marked A, B, or C indicate statistically significant differences (*p* < 0.05, two-way repeated measures ANOVA) in disease severity between mice fed a high-carbohydrate vs high-fat/high-protein diet (letter A), standard lab diet vs high-fat/high-protein diet (letter B), or high-carbohydrate vs high-fat/low-protein diet (letter C). There were significant (*p* ≤ 0.001, two-way repeated measures ANOVA, **) changes in CDI severity through time in infected mice fed a standard lab diet and a high-fat/high-protein diet. b) Kaplan-Meier survival curves for mice fed a high-carbohydrate diet (blue, n = 5), high-fat/low-protein diet (green, n = 5), high-fat/high-protein diet (red, n = 5), and standard lab diet (purple, n = 5), all following CDI challenge. The high-fat/high-protein diet significantly (*p* = 0.003, Log-Rank test, *) reduced survival of infected mice. All uninfected mice fed the standard lab diet showed no CDI signs (score of 0) for the duration of the experiment (not shown).

### Reduced microbial diversity is associated with changes in diet, antibiotic treatment, and CDI

To understand changes in gut microbial diversity due to experimental manipulations, alpha diversity was analyzed, as measured by richness (observed sequence variants (SVs), Fig. 3a), Simpson’s evenness (Fig. S1), and Shannon diversity (Fig. 3b) over the course of the experimental timeline. All animal groups showed significant (*p* < 0.05, ANOVA) change in diversity over time, due to decreases in richness and evenness corresponding to changes in diet (Day 13), antibiotic treatment (Day 17), and/or disease status (Day 18, 19).

**Fig. 3.**
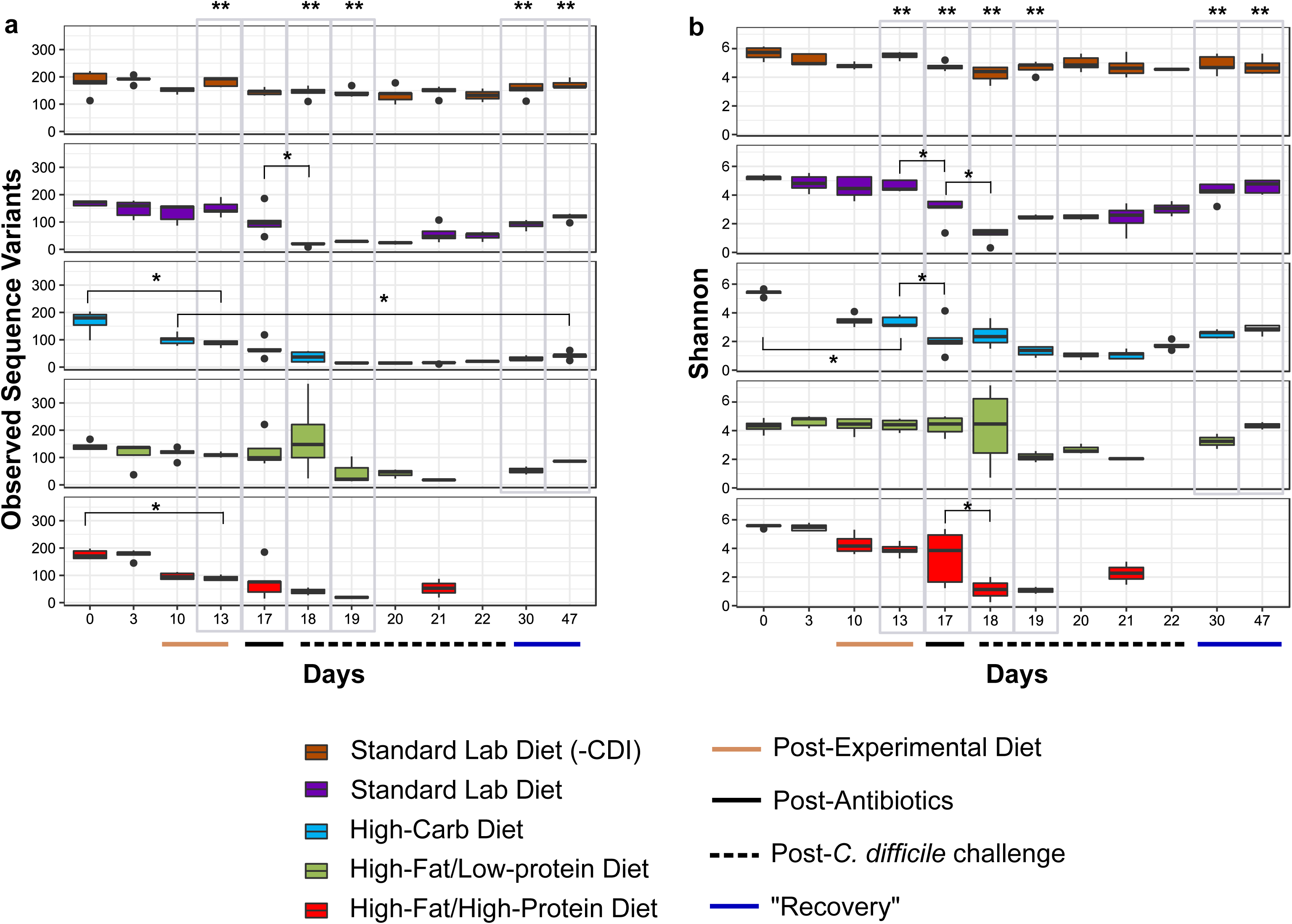
Effect of diet and treatment on alpha diversity. Observed sequence variants (SV) and Shannon diversity were calculated for uninfected mice fed a standard lab diet (orange, n = 5), and infected mice fed a standard lab diet (purple, n = 5), high-carbohydrate diet (blue, n = 5), high-fat/low-protein diet (green, n = 5), or high-fat/high-protein diet (red, n = 5). Gray boxes highlight comparisons between groups after a change in diet on Day 13, antibiotic treatments on Day 17, post-infection on Days 18 and 19, and recovery on Days 30 and 47. Administration of experimental diets (solid tan line, x-axis) and timepoints after antibiotics (solid black line, x-axis) and *C. difficile* challenge (dashed black line, x-axis) are indicated. Black dots above and below boxplots represent outliers. (*) Indicates significant (*p* < 0.05) loss of diversity in within-group pairwise comparisons. (**) indicate significant (*p* < 0.05) difference between groups on a given day (ANOVA).

Specifically, there was a significant difference (*p* < 0.05 ANOVA) in richness, evenness, and Shannon diversity between diet groups after mice were fed the experimental diets for ten days (Day 13), exemplified by significant decreases in richness and Shannon diversity after administration of the high-carbohydrate (Day 0 vs Day 13, *p* < 0.05, ANOVA and Tukey’s HSD). and richness after administration of the high-fat/high-protein diet (Day 0 vs Day 13, *p* < 0.05, ANOVA and Tukey’s HSD). Diversity was also distinct in the diet groups following antibiotic treatment; in particular, there was a significant loss of diversity after antibiotic treatments (Day 13 vs Day 17, *p* < 0.05, ANOVA and Tukey’s HSD) in mice fed the standard lab diet (evenness and Shannon diversity) and the high-carbohydrate diet (Shannon diversity). The diet groups were also distinct with regard to all three diversity indices following inoculation with spores (Day 18, Day 19), and there were significant losses in diversity in mice fed the standard lab diet and the high-fat/high-protein diet corresponding to CDI development (Day 17 vs Day 18, *p* < 0.05, ANOVA and Tukey’s HSD).

For mice fed the standard and high-fat/low-protein diets, most alpha diversity metrics returned to their diet-acclimated states within 30 days post-*C. difficile* challenge (Day 13 vs Day 47, *p* > 0.05). In contrast, gut microbiome richness did not return to normal in mice fed the high-carbohydrate diet over this time course (Day 10 vs Day 47, *p* < 0.05).

### Microbial communities transitioned through a common pattern in response to experimental manipulations

To assess microbial community changes over the experimental timeline, Bray-Curtis dissimilarity was calculated and visualized by NMDS. This analysis revealed a common pattern of microbial community transition through the experimental time course in all groups: diet-associated state, antibiotic-associated state (Abx), CDI-associated state (CDI), and recovery state (recovery) (Fig. 4). Although overall patterns in the progression of these states were similar, some diet-specific effects were apparent.

**Fig. 4.**
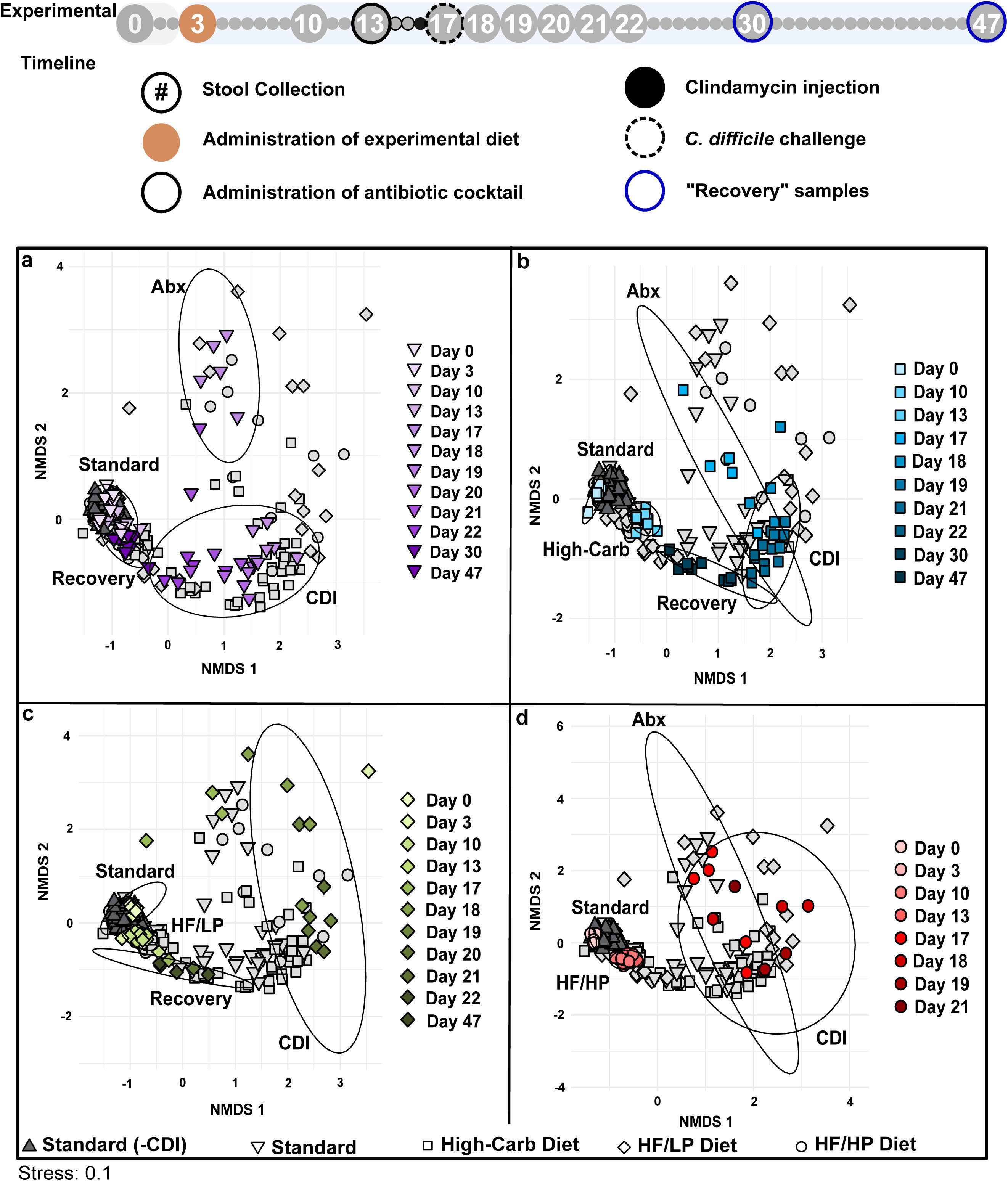
NMDS analysis based on Bray-Curtis dissimilarity. Each panel is a visualization of the same data and highlights the analysis for infected mice fed a A) standard lab diet, B) high-carbohydrate diet (blue), C) high-fat/low-protein diet (HF/LP, green), and D) high-fat/high-protein diet (HF/HP, red). Colors are shaded to show time progression through the experiment. Uninfected mice fed a standard lab diet (dark grey triangles) are featured in all panels. Ellipses represent standard errors of the mean (95% confidence) for samples associated with a the standard lab diet (labelled “Standard,” days 0 and 3), diet-associated microbiomes (days 10 and 13), antibiotic treatments (labelled “Abx,” day 17), CDI (days 18-22), and recovery (days 30 and 47) for colored data points associated with each experimental group are highlighted in the figure panel. A 95% confidence ellipse of samples representing mice fed a high-fat/low-protein diet on day 17 was not included, as it was large and included nearly all points in the data set. This indicates there is variability in these high-fat/low-protein samples after antibiotic treatments and results must be interpreted with caution due to small sample size. For guidance, an amended experimental timeline image from Fig. 1 is included.

Infected mice fed the standard lab diet progressed through distinct phases of transition and returned to a quasi-pre-CDI community structure, as indicated by overlapping “Standard” and “Recovery” confidence ellipses (Fig. 4a). However, some microbial groups did not return after the experimental treatments, exemplified by the phylum *Tenericutes* (Fig. S2–5). In contrast, the diet-associated and recovery ellipses did not overlap in mice fed the high-carbohydrate and high-fat/low-protein diets (Fig. 4b, 4c), indicating incomplete restoration of the gut microbial communities. The analysis also highlighted the large variability in microbial community structure in mice fed the high-fat/low-protein diet following antibiotic treatment (Fig. 4c), which was generally consistent with the heterogeneous CDI outcomes of the mice on that diet. A recovery phase was not observable in mice fed the high-fat/high-protein diet due to 100% mortality of these mice by day 21 (Fig. 4d). Uninfected mice fed the standard lab diet clustered together throughout the experiment.

### Diet impacted antibiotic- and recovery-associated microbial communities

To better investigate the effect of diet on microbial community response to specific treatments, Bray-Curtis dissimilarity was calculated for crucial time points in the experiment (Fig. 5). Diet-specific microbial communities developed prior to antibiotic treatment (Day 13), as indicated by non-overlapping ellipses and highly significant ANOSIM values (*p*<0.05; R>0.912) for standard, high-carbohydrate, and high-fat/high-protein diets on Day 13; however, the community structure of high-fat/low-protein-fed animals was not distinct (Fig. 5a).

**Fig. 5.**
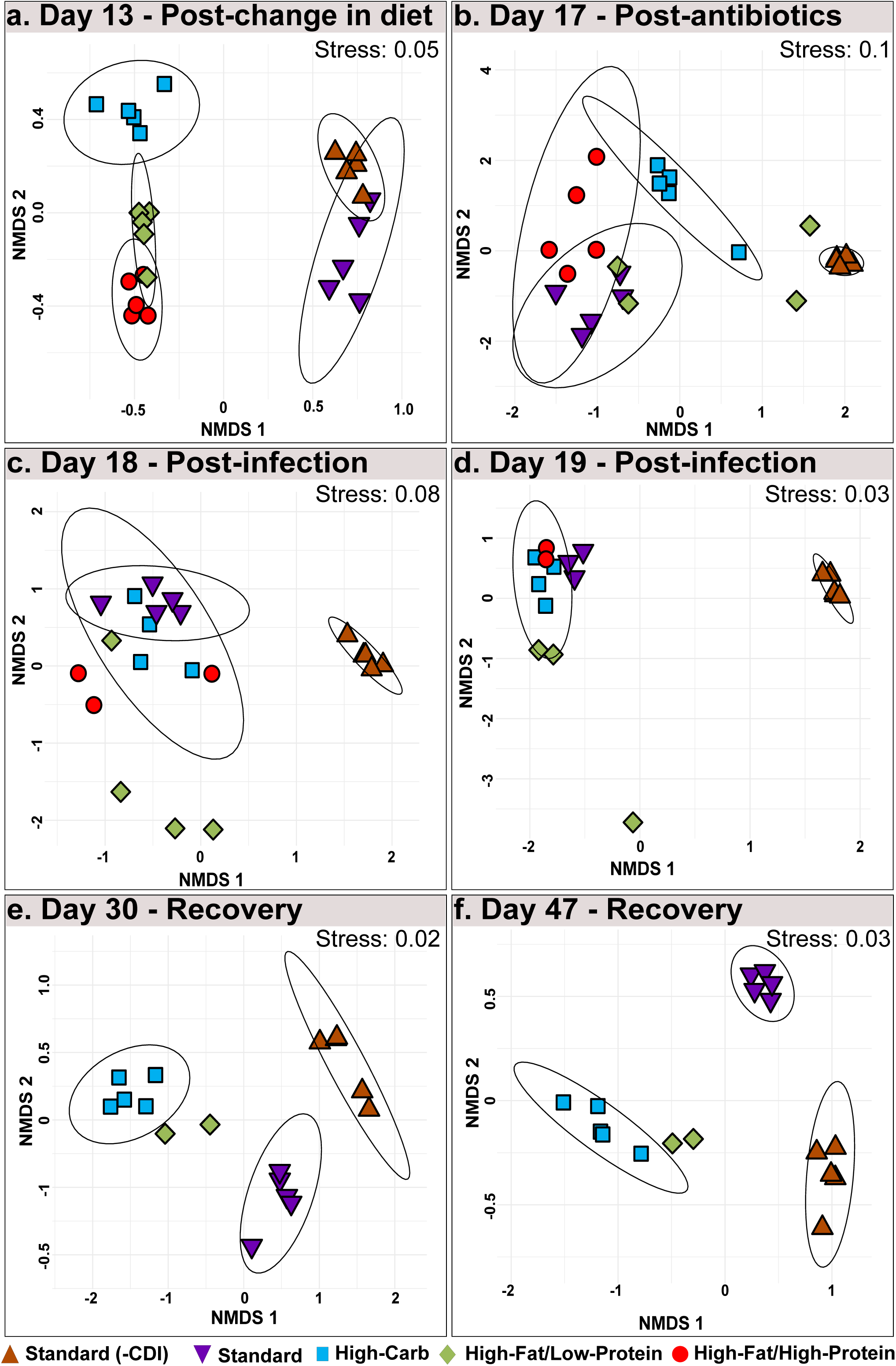
NMDS analysis based on Bray-Curtis dissimilarity for Days 13, 17, 18, 19, 30, and 47. Each panel is an ordination of samples from uninfected mice fed a standard lab diet (orange), and infected mice fed a standard lab diet (purple), high-carbohydrate diet (blue), high-fat/low-protein diet (green), or high-fat/high-protein diet (red) for A) Days 13, B) 17, C) 18, D) 19, E) 30, F) and 47. Ellipses represent standard errors of the mean (95% confidence). 95% confidence are not shown mice fed a standard lab diet and high-fat/low-protein diet on day 17, 18, and 19, as they were large and included nearly all points in the data set. 95% confidence ellipses are not shown for mouse groups with significant mortality, as the calculation cannot compute with n < 4.

The distinctness of the microbial communities was disrupted by antibiotic treatment and CDI, as evidenced by overlapping ellipses and insignificant ANOSIM values for Day 17, Day 18, and Day 19, indicating the dominant role of the antibiotic treatments and CDI over the diet treatments in structuring the microbial community (Fig. 5b-d). Following recovery, diet-specific clustering patterns re-emerged in the recovery phase on Day 30 and Day 47 (Fig. 5e-f).

Similar to CDI severity signs, the high-fat/low-protein microbiomes were heterogeneous during these treatments. However, no connection was observed between individual animal changes in microbial communities and disease severity or onset.

### Diet and antibiotics administration profoundly altered the microbiome composition

SIMilarity PERcent (SIMPER) analysis (22) identified 51 SVs that contributed to 50% of microbial community dissimilarity between all pairwise comparisons of the diet-specific microbiomes throughout the experiment (Fig. 6). More than half of these SVs belonged to the *Clostridiales*, predominantly the families *Lachnospiraceae* (19/51) and *Ruminococcaceae* (9/51) and were dominated by uncultivated genera. Most *Lachnospiraceae* SVs decreased in abundance after administration of the experimental diets, particularly the high-fat/low-protein diet and the high-carbohydrate diet, and were further reduced following antibiotic treatment and CDI. The *Ruminococcaceae* SVs were more variable in response to the diets but were also strongly depleted following antibiotic treatment.

**Fig. 6.**
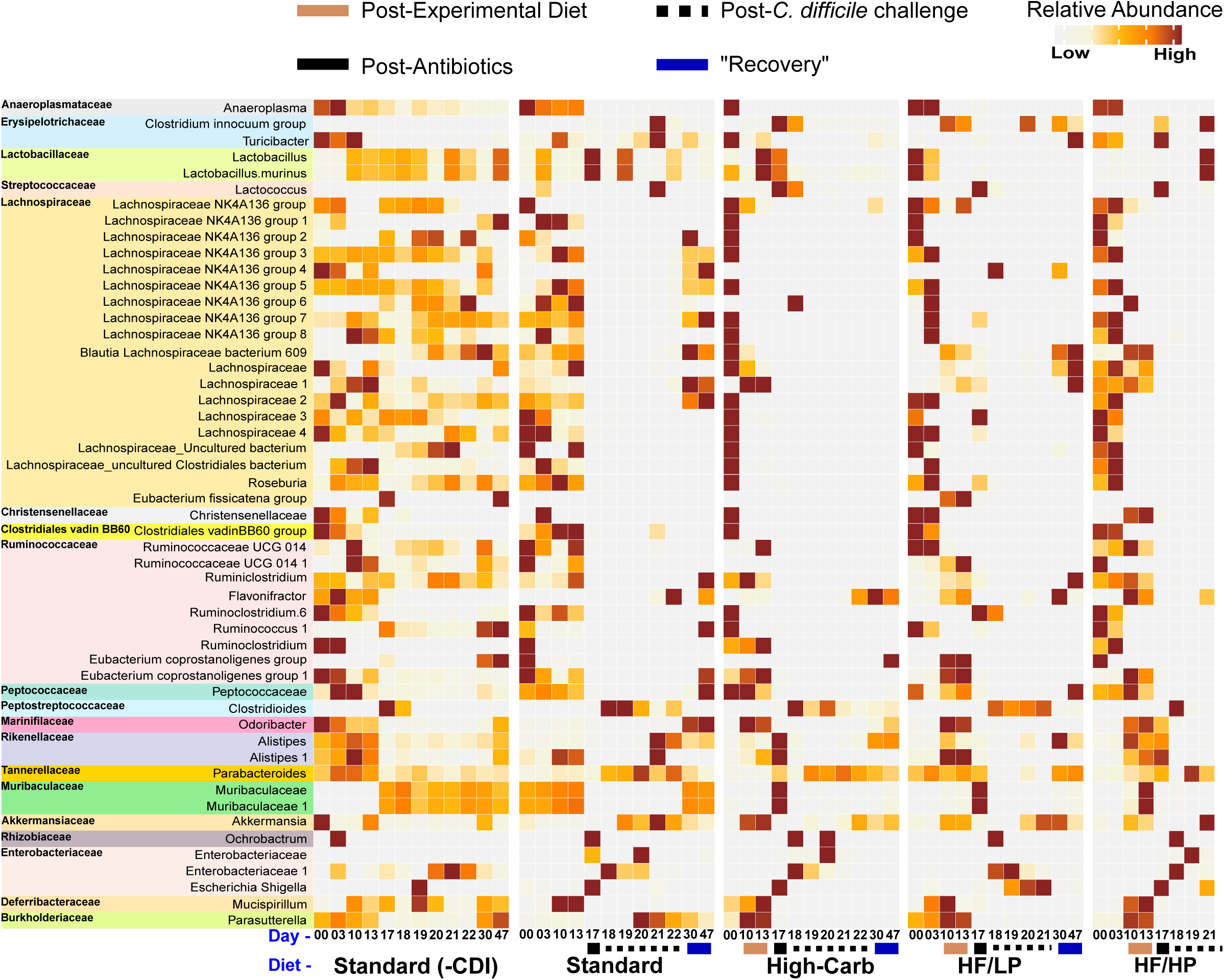
SIMPER analysis results displaying top SVs responsible for dissimilarity between experimental groups. Heatmap of the mean relative abundance of 51 SVs that contributed cumulatively to 50% of community dissimilarities at each time point among the standard lab diet, high-carbohydrate diet (High-Carb), high-fat/low-protein diet (HF/LP), and high-fat/high-protein diet (HF/HP). Each square represents the mean relative abundance of the given SV on a particular day for a particular diet. Higher intensity of the brown color correlates with high relative abundance.

Several other groups also showed strong patterns. Two *Muribaculaceae* SVs became depleted in the high-carbohydrate and high-fat/low-protein groups but then bloomed and crashed following antibiotic treatment and never recovered. Also, two *Alistipes* SVs slightly increased after administration of the experimental diets, followed by a reduction after antibiotic treatment and/or CDI. An SV affiliated with the *Clostridium innocuum* group emerged at different times after the antibiotic treatment in all the antibiotic-treated mice. Further, some members of the *Proteobacteria*, including *Escherichia*/*Shigella* and an uncultivated member of the *Enterobacteriaceae*, expanded after the antibiotic treatment, yet *Parasutterella* decreased in all antibiotic-treated mice. Also, an SV in the order *Parabacteroides* expanded after the antibiotic treatment in all but high-fat/low-protein diet treatments. The relative abundance of *Akkermansia* increased after *C. difficile* challenge (Day 21) irrespective of diet.

## Discussion

The mammalian gut microbiota is crucial for host health, and also provides colonization resistance against various enteric pathogens (23). Exposure to broad-spectrum antibiotics leads to the depletion of commensal microbiota, which can be exploited by pathogens such as *C. difficile* (24). Diet is an important force that determines gut microbial composition and function (25). Consequently, several studies have demonstrated the effects of dietary components on *C. difficile* growth, physiology, and pathogenesis both *in vitro* and in antibiotic-induced animal models of CDI; however, those studies are contradictory with regard to the relative importance of proteins and carbohydrates in CDI. The goal of this study was to broadly assess CDI outcomes and microbial community responses to diets with extreme differences in macronutrient composition following antibiotic treatment.

### Effect of high-carbohydrate diets on antibiotic-induced CDI

Improved gut health due to high-carbohydrate diets, especially those rich in fiber, has been well-documented and is suggested to be related to the production of SCFA by gut microbes (26–29). Some studies point specifically to the importance of MACs in mitigation of CDI, specifically inulin (10); however, the inulin content of the high-carbohydrate diet in our study was low (2.1% w/v). Instead, the major sources of carbohydrates were corn starch (43.5% w/v), maltodextrin (14.4% w/v), and sucrose (11.0% w/v). the highly digestible corn starch and maltodextrin incorporated in these diets would be depolymerized to monosaccharides readily. Thus, our study suggests that a high-carbohydrate diet that is correspondingly low in protein can be protective against CDI, irrespective of the specific carbohydrate composition.

This result is superficially contradictory to reports of recent adaptations enhancing the transport, metabolism, and general physiology of different strains of *C. difficile* in response to glucose, fructose, and trehalose (14, 15). However, the study on glucose and fructose metabolism did not describe any effects of sugars on pathogenesis of RT027 and it is possible that some substrates (e.g., monosaccharides) promote the maintenance and spread (via enhanced spore production and survival) of *C. difficile*, while simultaneously limiting *C. difficile* overgrowth. Indeed, we found that mice fed the high-carbohydrate diet became long-term carriers of low-abundance populations of *C. difficile*, whereas mice fed the standard and high-fat/low-protein diets cleared *C. difficile* within 30 days post-*C. difficile* challenge (Fig. S6). These studies paint a somewhat complicated picture for the effect of dietary sugars on CDI that should be addressed using carefully controlled and documented experiments that take into account the strain of *C. difficile* and the precise diet used in the animal model. In this regard, it is noteworthy that recent studies on the effect of sugars on *C. difficile* do not report the diet used for the experiments (14, 15).

### Effect of a high-fat/high-protein, Atkins-like diet on antibiotic-induced CDI

Our experiments revealed that CDI was exacerbated in mice fed high-fat diets, particularly in mice fed a high-fat/high-protein, Atkins-like diet. A high-fat diet was also observed to intensify CDI in a hamster model of infection (30); however, to our knowledge, a high-fat/high-protein diet has never been explored in animal models of CDI, as typical diets used for mouse and hamster models of CDI have ≤ 10% kcal from fat and ≤ 30% kcal from protein. The poor CDI outcome observed here for mice fed the high-fat/high-protein diet compared with the high-fat/low-protein diet is consistent with known relationships between dietary protein and CDI in both lab mice with native murine microbiota (13) and humanized mice fed a defined amino acid diet deficient in the Stickland electron acceptor proline (11). The major diets used in these other studies used relatively low concentrations of protein (≤ 20%) provided as casein. The protein source in our high-fat/low-protein diet was also casein (22.8% w/v, File S2), yet that for the high-fat/high-protein diet was whey protein (50.0% w/v, File S1) (31). However, since both casein and whey protein are derived from milk and have similar amino acid content (File S1 and 3), and they are provided as pre-hydrolyzed peptides, we expect that they will differ little in amino acid availability. However, because the protein sources were not identical, further work is needed to disentangle any effects of protein source, digestibility, or abundance on CDI.

Effects of protein on CDI could be direct and/or indirect. *C. difficile* can use many different amino acids as Stickland donors and both proline and glycine can be used as Stickland acceptors (9). Yet, the *C. difficile* proline reductase, PrdA, was only expressed in humanized mice inoculated with dysbiotic microbiomes (11), suggesting *C. difficile* may not compete well for proteins or amino acids with healthy microflora. In “healthy” humanized mice, PrdA was expressed by three members of the *Lachnospiraceae*, suggesting they might be important competitors for proline and other amino acids. *Lachnospiraceae* was also one of only three bacterial families that was significantly depleted in the dysbiotic humanized mice, along with *Ruminococcaceae* and *Bacteroidaceae*. In our study, the majority of the bacteria that contributed to microbiome variation belonged to *Lachnospiraceae* or *Ruminococcaceae*, and all of these organisms decreased in abundance in response to the diets and/or antibiotic treatment, suggesting they may be important competitors of *C. difficile* for amino acids in the lumen (32). *Lachnospiraceae* in particular is a dominant bacterial family in the gut microbial communities of many mammals and many provide benefits to the host (33). A member of this family has also been documented to provide colonization resistance to *C. difficile* in a CDI mouse model (34), and have been shown to produce iso (3β-hydroxy) bile acids (35), which are inhibitors of *C. difficile* growth. Thus, it appears that members of *Lachnospiraceae* and *Ruminococcaceae* act against *C. difficile* through independent mechanisms, including competition for resources and production of secondary bile acids. In our study, the high-fat/high-protein diet would lead to an abundance of oligopeptides and free amino acids in the lumen and could provide a selective advantage that leads to *C. difficile* overgrowth when coupled with the loss of *Lachnospiraceae* and *Ruminococcaceae.* Diet or antibiotic-induced loss of *Lachnospiraceae* and *Ruminococcaceae* might not be a significant problem in mice fed other diets, particularly the high-carbohydrate diet, because of the low concentrations of peptides and amino acids and/or the healthful effects of SCFAs resulting from carbohydrate fermentation by other microorganisms (see above).

High-protein diets have more general consequences for gut health that may exacerbate pathogenesis. Proteins that are not easily digested causes a surplus of dietary protein reaching the distal gut that can be acted upon by protein-fermenting bacteria. Fermentation of sulfur-containing amino acids can lead to excessive hydrogen sulfide, via desulfurlyation of cysteine and methionine, which inhibits host cytochrome c oxidase, damages colonic cell genomic DNA, alters cellular pathways, inhibits host-cell butyrate oxidation, and reduces the lifespan of colonocytes (36, 37). Production of ammonia due to protein fermentation (ammonification) can result in reduced gut health through decreased SCFA production, increased paracellular permeability of gut epithelial cells, and altered epithelial cell morphology (38, 39). Yet, the effect of the high-fat/high-protein diet on mice health alone does not explain the health problems in those mice, as the experimental diets were given for ten days prior administration of antibiotics and *C. difficile* challenge and there were no observable changes in behavior or host health.

### Effect of a high-fat/low-protein keto-like diet on antibiotic-induced CDI

Mice fed the high-fat/low-protein diet showed variability in disease severity and survival (Fig. 2) and highly individual variable microbiome composition after antibiotic treatments (Fig. 4). Thus, the effect of fats on CDI is uncertain. The high-fat/low-protein diet was high in saturated fats, as the fat source was hydrogenated coconut oil (33.3% w/v, File S2). Coconut oil is >90% saturated fatty acids and consists of medium-length chains rich in lauric acid (C8:0) (40). This diet closely mimics the MCT diet, which is a version of ketogenic diet (19). Diets high in saturated fats can increase incidence of colitis in immunocompromised mice due to increased inflammatory response (41). This points to plausible mechanisms by which healthy mice fed the high-fat/low-protein diet may have experienced increased CDI severity and lower survival post-challenge. However, the high-fat diets used in these studies also had low carbohydrate content, which itself may have been an important factor in disease progression in some animals.

### Other microbial community responses to diets, antibiotics, and CDI

Following changes in diet, mice developed distinct microbial communities, which were then disturbed by antibiotic treatments and CDI (Fig. S2–5, Fig. 4). The shift in community composition after diet and antibiotics was accompanied by reduced alpha diversity after diet, antibiotic treatment, and CDI (Fig. 3 and Fig. S1), which is consistent with previous work. Perturbations to gut microbial communities from antibiotics can result in CDI due to loss of colonization resistance, especially following a course of antibiotics, as seen in humans (42) and across CDI mouse and hamster animal models (5, 43, 44). Typically, loss of colonization resistance and susceptibility to CDI is marked by a decrease in gut microbial diversity (5, 45), yet a recent report suggests CDI susceptibility and severity are independent of alpha diversity (10). Likewise, our work shows loss of diversity in infected mice across all experimental diets (Fig. 3), yet significantly different survival and severity outcomes between the groups (Fig. 2). This demonstrates that diet plays a critical, and dominant, role in CDI disease patterns to microbial diversity *per se*.

In addition to changes in *Lachnospiraceae* and *Ruminococcaceae*, described above, several other microorganisms changed throughout the experiment. There was a loss of a member of the genus *Parasutterella* after a change in diet (Fig. 6). *Parasutterella* species are a part of the core gut microbiome in humans and mice with occasional associations with disease. A study characterizing *Parasutterella* isolated from mouse guts showed that this organism significantly changed aromatic amino acid metabolism in the gut and is crucial for bile acid metabolism and homeostasis (46). This is especially important since conjugated primary bile salts are required for spore germination while secondary bile acids are known inhibitors of sporulation and growth of *C. difficile*. Therefore, *Parasutterella* could play an important role in colonization resistance through the modulation of the composition of the bile acids pool (47).

*Muribaculaceae* (formerly Family S24-7) decreased after administration of the high-carbohydrate and high-fat/low-protein diets (Fig. 6). *Muribaculaceae* is well-adapted to the mouse gut microbiome (48), and may contain up to 685 species-level clusters (49). Further, analysis of the 157 draft genomes of the members of *Muribaculaceae* showed them to be highly enriched in glycoside hydrolases, suggesting an ability to deconstruct complex carbohydrates such as MACs and thereby promote SCFA production (49, 50).

The increase in *Alistipes* abundance after the diet changes and their fall after antibiotic treatment could be indicative of general antibiotic susceptibility of this organism. Members of *Alistipes* are suggested to be protective against CDI (44) and depletion of these microbes has been associated with CDI (51). Another study on fecal microbiota transplant (FMT) treatments for CDI documented an increased abundance of *Alistipes* after the FMT (52). Hence, the protective effect provided by members of the genus *Alistipes* needs to be evaluated carefully.

The increased abundance of *Clostridium innocuum* after antibiotic treatment in all diet groups could be attributed to its ability to produce peptidoglycan precursors with a C-terminal serine, which have a low affinity for vancomycin (53). This organism’s resistance to vancomycin is a potential cause of antibiotic-associated diarrhea in humans similar to CDI (54).

Increased *Akkermansia* post-*C. difficile* infection could be linked with increased mucus levels. In human CDI, patients with active CDI tend to secrete acidic mucins composed of MUC1 with altered oligosaccharide composition (55). We speculate that the increased abundance of *Akkermansia* post-infection (Fig. 6) could be a result of this mucus secretion (56).

The transient surge in *Proteobacteria* after antibiotic treatment in all mice groups is a known response to loss of the dominant gut taxa (57). *Proteobacteria* induce inflammation (57) and provide a conducive environment for invasion by pathogens like *C. difficile* (58). However, this increase in members of the *Proteobacteria* was also seen in mice fed the high-carbohydrate diet, indicating that blooms of *Proteobacteria* are insufficient to induce CDI. *Parabacteroides* also increased in patients with recurrent CDIs (59). *Parabacteroides* produces succinate (60), which has been found to promote *C. difficile* growth following antibiotic treatments (61). Thus, the survival of *Parabacteroides* after antibiotic treatments could favor the proliferation of *C. difficile* in infected mice.

## Conclusion

This study shows large differences in the outcome of CDI in an antibiotic-induced mouse model using hypervirulent strain R20291 (RT027) due to diets representing extremes of macronutrient composition. The poor outcome of mice fed an Atkins-like diet demands a close look at whether Atkins, ketogenic, or other high-fat/high-protein diets create high risk for CDI in humans, particularly if members of the *Lachnospiraceae* and *Ruminococcaceae* are disrupted by antibiotics. In contrast, the protective effect of a high-carbohydrate diet, despite high monosaccharide and digestible starch content is inconsistent with recent reports on the possible importance of simple sugars on CDI. Since monosaccharides and digestible starch would not be expected to improve gut health, the apparent protective effect may be due to the low protein and/or fat content rather than protective effects of carbohydrates *per se*.

## Methods

### Bacterial growth conditions and spore harvest

*C. difficile* R20291 (RT027), a gift from Nigel Minton, University of Nottingham, was grown on Bacto BHI agar plates in an anaerobic chamber (10% CO_2_, 10% H_2_, 80% N_2_) at 37°C for 7 days. Bacterial cells and spores were collected from plates with confluent growth by flooding with ice-cold, autoclaved deionized water. Spores were pelleted by centrifugation and resuspended in fresh deionized water for three wash steps. The spores were harvested by density-gradient centrifugation via a 20% to 50% HistoDenz gradient. The spore pellet was washed five times with autoclaved deionized water and stored at 4°C. Schaeffer-Fulton-staining was performed to determine purity of spore harvest.

### Animals

The animal protocol 1039564-2 used in this study was approved by the Institutional Animal Care and Use Committee (IACUC) at the University of Nevada, Las Vegas. All experiments and procedures were conducted in line with the National Institutes of Health’s guidelines in the Guide for Care and Use of Laboratory Animals. Bedding, water, and mice feed were autoclaved prior to use. Weaned female C57BL/6 mice were purchased from Charles River (Wilmington, MA) and given one week to acclimate in the animal facility. The following protocols represent animals 5-8 weeks old.

### Diet specifications

The mice chow used for each experimental diet (File S1–3; Table S1) were purchased from TestDiet and irradiated prior to shipping. The mice chow was stored at 4°C before use. Additional diet specifications can be found in Files S1–3.

### Treatment groups and experimental timeline

Mice were caged in groups of five and each mouse was marked with one of five colors. Feces was collected from each individual mouse by stroking the back of the mouse to induce defecation. Fecal samples were collected beginning at Day 0 and for the duration of the 47-day experiment. All animals were fed a standard lab diet until Day 3. Animals were randomly separated into 5 groups (Fig. 1, Table S1). Two groups received a standard lab diet (Standard lab diet and untreated controls Standard (-CDI)). Three groups received a high-carbohydrate, high-fat/low-protein, or a high-fat/high-protein diet. All groups were fed their respective diet and autoclaved water ad libitum. On Day 13 and for the next two days, all groups except Standard (- CDI) were given an antibiotic cocktail (ad libitum in sterile drinking water, changed daily) containing kanamycin (0.4 mg/mL), gentamicin (0.035 mg/mL), colistin (850 U/mL), metronidazole (0.215 mg/mL), and vancomycin (0.045 mg/mL). After antibiotic treatment, all animals were given regular drinking water for the remainder of the experiment. On Day 16, all groups except Standard (-CDI) were administered a single dose of intraperitoneal clindamycin (10 mg/kg) dissolved in autoclaved DI water. Animals in Standard (-CDI) were administered intraperitoneal DI water, the vehicle control for clindamycin. On Day 17, animals in all groups, except standard lab diet -CDI, were challenged with 10^8^ CFUs of *C. difficile* R20291 spores by oral gavage. For 30 days following challenge, animals were monitored daily for signs of CDI. CDI severity were scored following the rubric described in the section below. Mice were weighed twice daily after challenge inside a biosafety hood. Animals were euthanized 30 days post infection, or as soon as CDI symptoms reach a clinical endpoint described below, so that no animals experienced unrelieved pain or distress.

### Scoring CDI severity

For mice, disease signs were scored using the following rubric, amended from previously published protocols (21): pink anogenital area (score of 1), red anogenital area (score of 2), lethargy (score of 2), diarrhea/increase in soiled bedding (score of 1), wet tail (score of 2), hunchback posture (score of 2), 8-15% loss of body weight (score of 1), >15% loss of body weight (score of 2). Animals scoring 2 or less were indistinguishable from non-infected controls and were considered non-diseased. Animals scoring 3–4 were considered to have mild CDI with signs consisting of pink anogenital area, lethargy, an increase of soiled bedding and minor weight loss. Animals scoring 5–6 were considered to have moderate CDI with signs consisting of mild CDI signs plus red anogenital area and hunchback posture. Animals scoring >6 were considered to have severe CDI and were immediately euthanized. Dietary changes were not expected to affect the animal negatively. However, if any signs of distress, or disease were observed, procedures were carried out as indicated in the rubric. A two-way repeated measures ANOVA was calculated to determine the effect of different diets over time on disease severity. Euthanized mice were also included in the ANOVA calculation and were given a score of 7. Statistical significance was defined as (p < 0.05).

### Illumina sequencing

Fecal samples for microbiome analysis were collected and archived at -80°C. DNA was extracted from thawed fecal samples using the QIAamp DNA Stool Mini Kit and quantified using a NanoDrop 1000 Spectrophotometer. The V4 region of the 16S rRNA gene was PCR amplified using modified primers 515F:GTGYCAGCMGCCGCGGTAA and 806R: GGACTACNVGGGTWTCTAAT, with the forward primer containing 12 bp barcode (62). The PCR products were cleaned, quantified and pooled at equimolar concentration. Paired-end sequencing (151bp x 12bp x 151bp) of pooled amplicons was performed using MiSeq platform and customized sequencing primers at Argonne National Laboratory. Over 6.4 million good quality reads of V4 region of 16S rRNA gene from 237 fecal samples were collected throughout the experimental timeline.

### Sequence analysis

All sequence-based analysis was performed in QIIME 2 (version 2018.6) (63). Raw Illumina reads were demultiplexed using sample-specific barcodes; approximately 29146 reads per sample were obtained. The sequences were denoised using dada2-denoise-paired plugin to remove low quality, chimeric, and artifact sequences and the resulting high-quality sequences were clustered into 2657 sequence variants (SVs). Taxonomy was assigned to each SV using sklearn-based taxonomy classifier that uses a Naive Bayes machine learning for classification and classifier that was trained on V4 region of 16S rRNA gene ‘Silva 132 99% OTUs full-length sequences’ database. SVs assigned to mitochondria, chloroplast, eukarya, and other unknown domains were excluded from further analysis. Phylogenetic tree of SVs was created by performing multiple sequence alignment using MAFFT followed by constructing an unrooted tree using FastTree which was then rooted using midpoint rooting method. The resultant tree was utilized for subsequent phylogeny-based analysis.

### Analysis of the effects of diet on the gut microbiome

Alpha diversity was estimated by calculating Shannon, Simpson, and Observed SVs diversity indices. An ANOVA test was performed to test if there were significant differences in the alpha diversity in response to diet, antibiotic administration, and CDI. Further, a Tukey’s honest significance test on the ANOVA output was performed to conduct pairwise comparisons of alpha diversities indices and correct for multiple comparisons. The data includes missing data due to mice death, or inability to collect fecal samples due disease severity. SV table, phylogenetic tree, and associated metadata file were then imported in R (3.5.0) (64) and further analyzed using phyloseq (version 1.25.2) (65), and vegan (version 2.5.2) and suitably visualized using ggplot2 (version 3.0.0) and Inkscape. NMDS was performed using Bray-Curtis dissimilarity and the relationship of the samples with respect to diet, antibiotic treatment, and CDI over the course of experiment was displayed using the first two dimensions. ANOSIM was calculated using a Bray-Curtis dissimilarity matrix. SIMPER was used to identify the SVs that are contributing to 50% of the observed differences in microbial communities at any time point during the experiment.

### Data Availability

Files containing the original unfiltered sequences are available from the NCBI-SRA under accession number SRP188890.

## Acknowledgements

This research is supported by the NIH under grant 1R01AI109139-01A1 and a UNLV Faculty Opportunity Award.

## Author information

### Contribution

E. A-S., and B. P. H. conceived and designed the experiments. C. C. M, J. R. P, J. V. V., S. A., and D. M. D. conducted animal experiment and collected samples. C. C. M, J. V. V., and S.A. processed the samples. S. S. B. carried out bioinformatic and statistical analysis, C. C. M., S. S. B., and B. P. H. interpreted the data. C. C. M, S. S. B, E. A-S, and B. P. H. wrote the manuscript. E. A-S. and B. P. H. were responsible for funding acquisition. All authors read, edited, and approved the final manuscript.

### Competing interests

The authors declare no competing interests.

## Supplemental figure legends

**Fig. S1.**
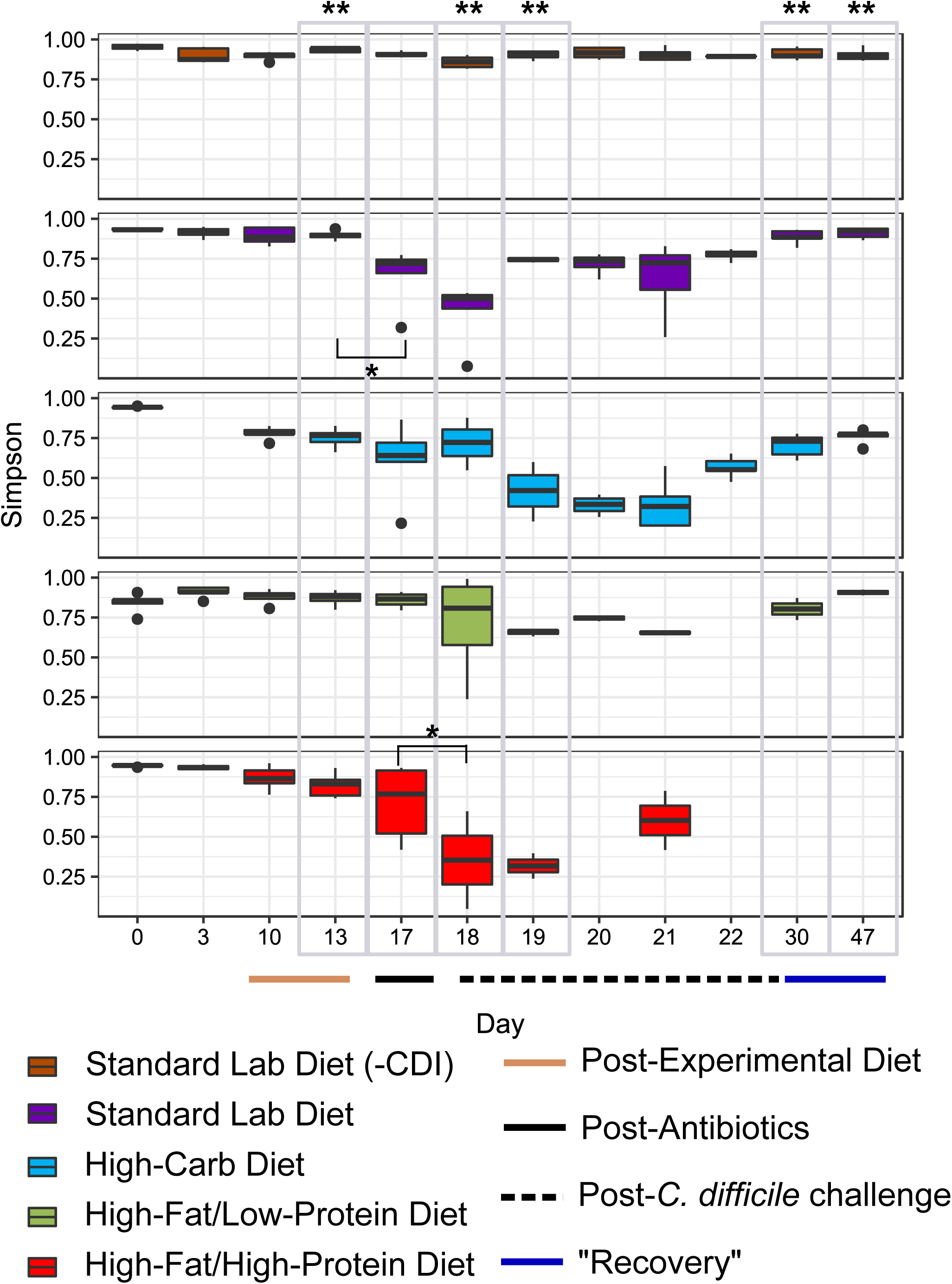
Effect of diet and treatment on alpha diversity – Simpson’s Evenness. Simpson’s evenness was calculated for uninfected mice fed a standard lab diet (orange, n = 5), and infected mice fed a standard lab diet (purple, n = 5), high-carbohydrate diet (blue, n = 5), high-fat/low-protein diet (green, n = 5), or high-fat/high-protein diet (red, n = 5). Gray boxes highlight comparisons between groups on after a change in diet on Day 13, antibiotic treatments on Day 17, post-infection on Days 18 and 19, and recovery on Days 30 and 47. Administration of experimental diets (tan circle, x-axis) and timepoints after antibiotics (gray circle, x-axis) and C. difficile challenge (dashed circle, x-axis) are indicated. (*) indicates significant (p < 0.05) loss of diversity in within group pairwise comparisons. Significance was calculated using a two-way repeated measures ANOVA.

**Fig. S2.**
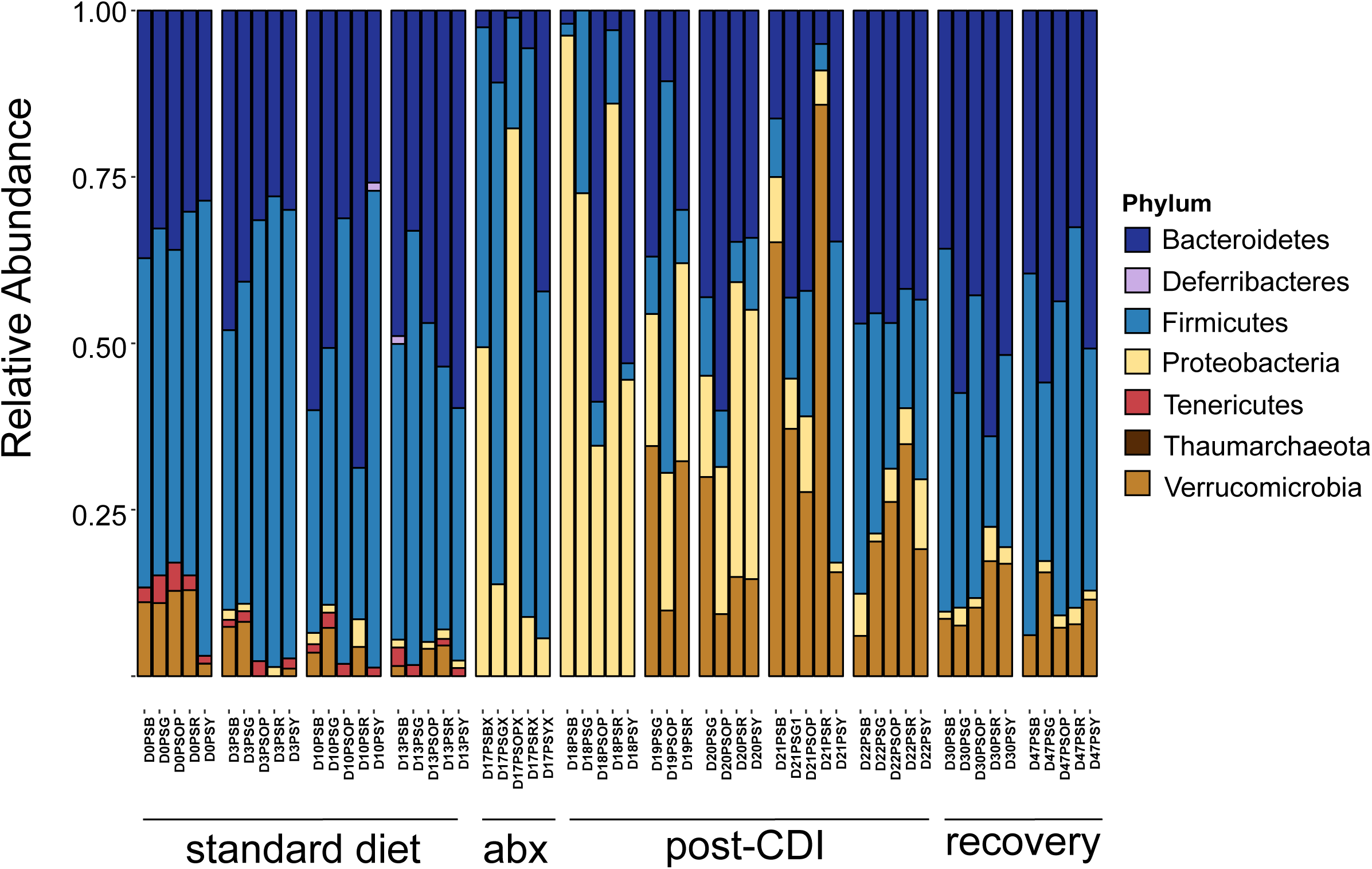
Relative abundance bar plot – standard lab diet. Bar plot showing the relative abundance of gut microbiota at the phylum level in each mouse fed the standard lab diet across the experimental timeline. Samples associated with days mice were fed a standard lab diet, administered antibiotics, post infection, and during recovery are indicated.

**Fig. S3.**
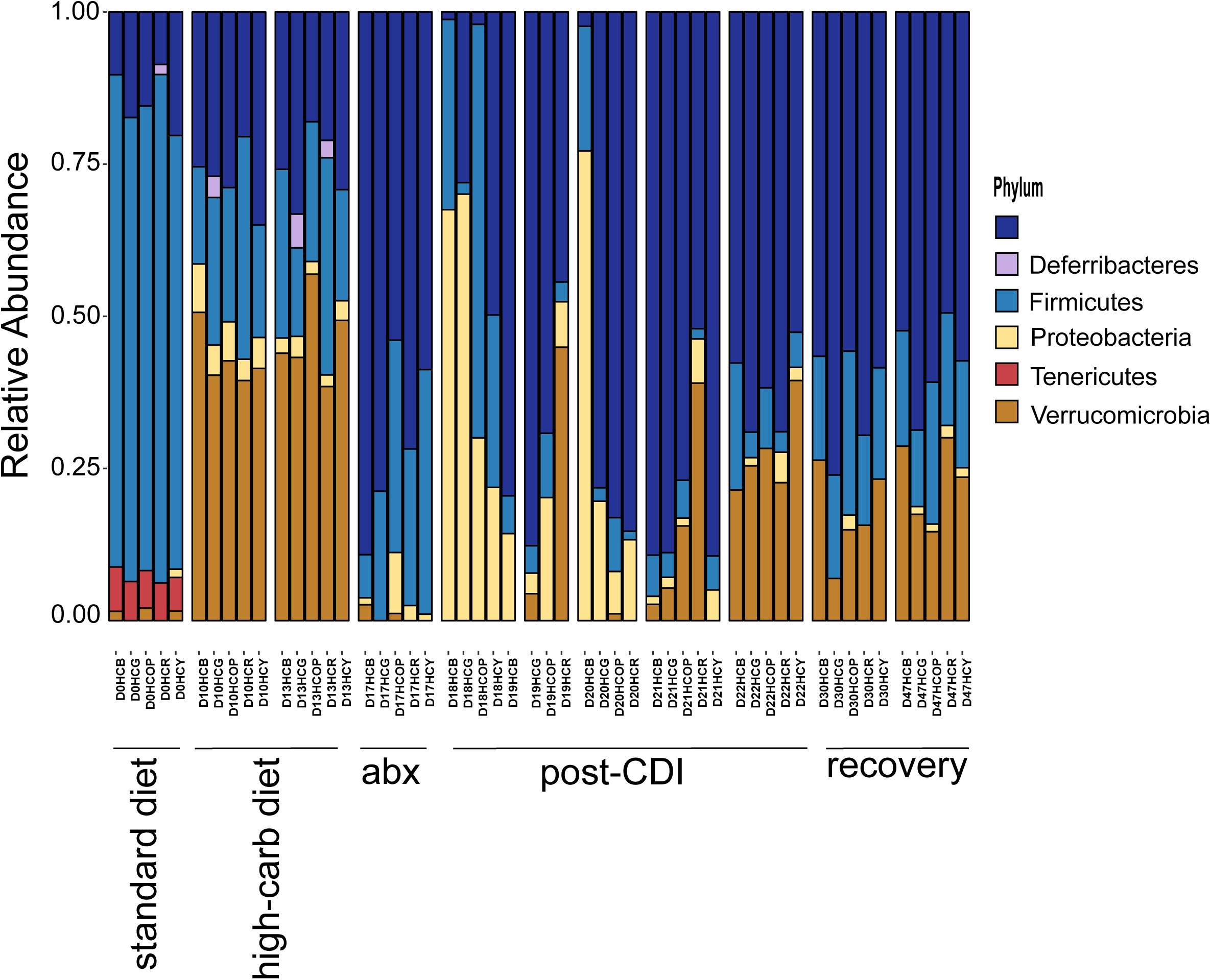
Relative abundance bar plot – high-carbohydrate diet. Bar plot showing the relative abundance of gut microbiota at the phylum level in each mouse given the high-carbohydrate diet across the experimental timeline. Samples associated with days mice were fed a standard lab diet, switched to the high-carbohydrate diet, administered antibiotics, post-infection, and during recovery are indicated.

**Fig. S4.**
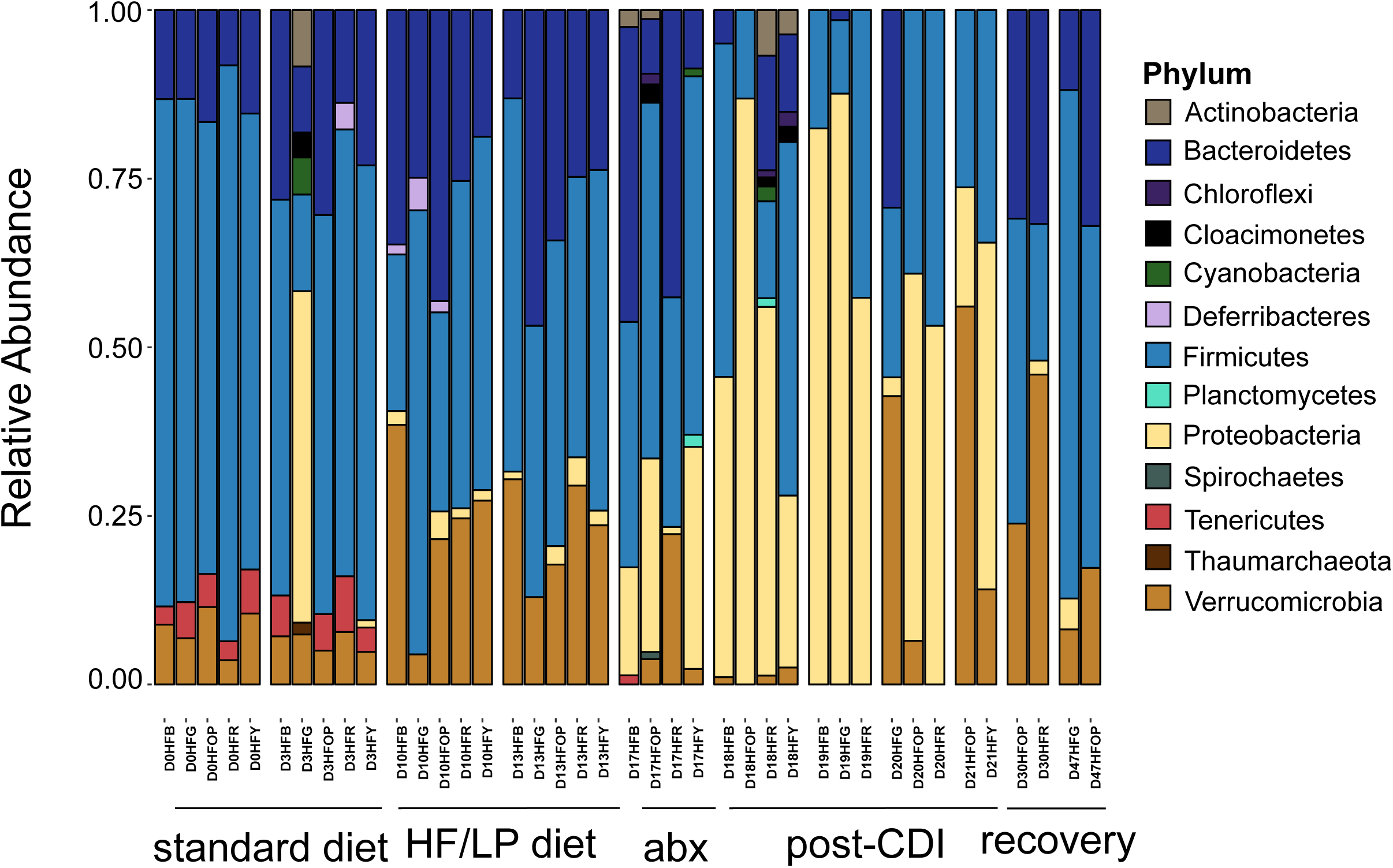
Relative abundance bar plot – high-fat/low-protein (HF/LP) diet. Bar plot showing the relative abundance of gut microbiota at the phylum level in each mouse given the high-fat/low-protein (HF/LP) diet across the experimental timeline. Samples associated with days mice were fed a standard lab diet, switched to the high-fat/low-protein diet, administered antibiotics, post-infection, and during recovery are indicated.

**Fig. S5.**
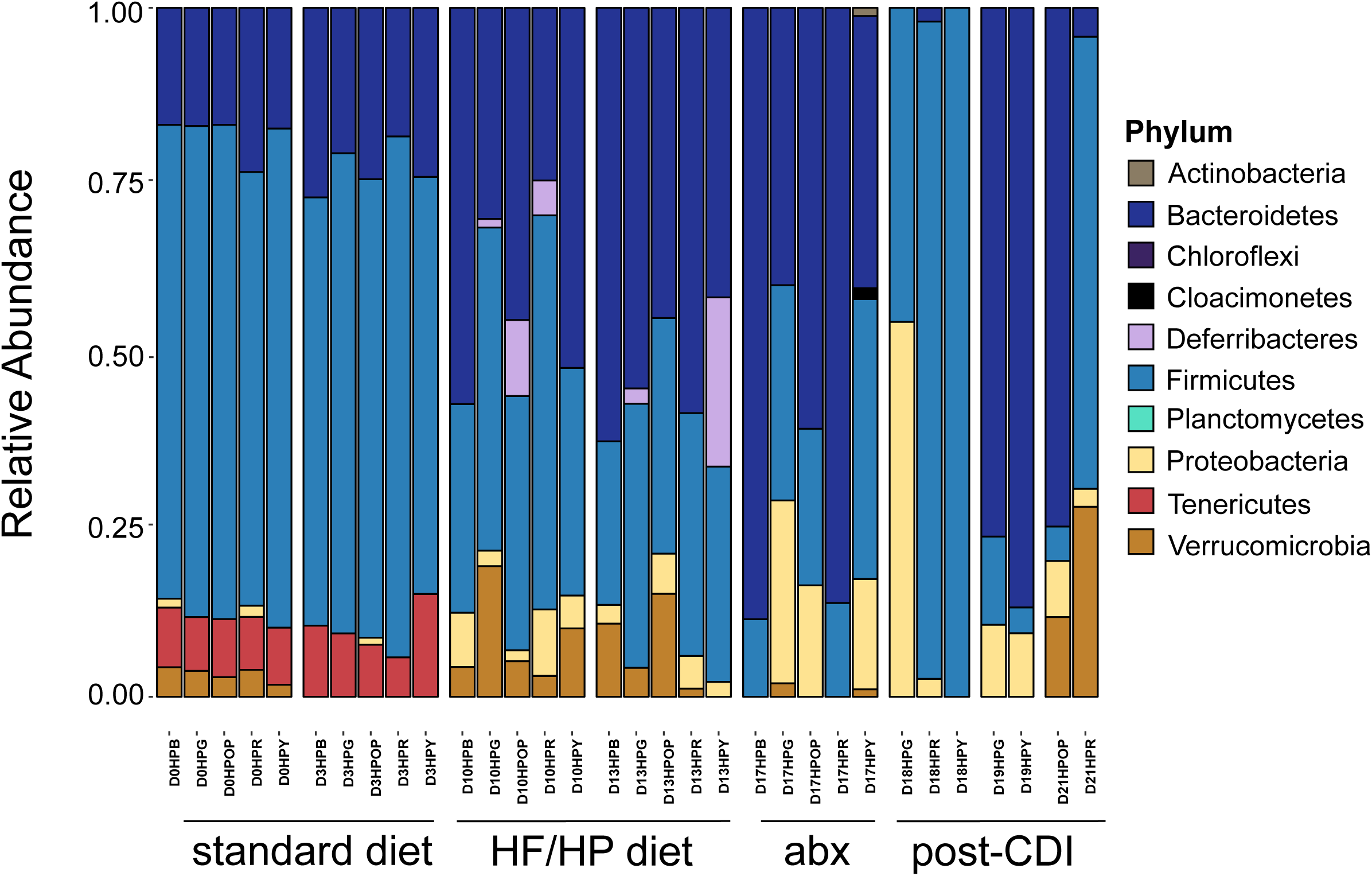
Relative abundance bar plot – high-fat/high-protein (HF/HP) diet. Bar plot showing the relative abundance of gut microbiota at the phylum level in each mouse given the high-fat/high-protein (HF/HP) diet across the experimental timeline. Samples associated with days mice were fed a standard lab diet, switched to the high-fat/high-protein diet, administered antibiotics, post-infection, and during recovery are indicated.

**Fig. S6.**
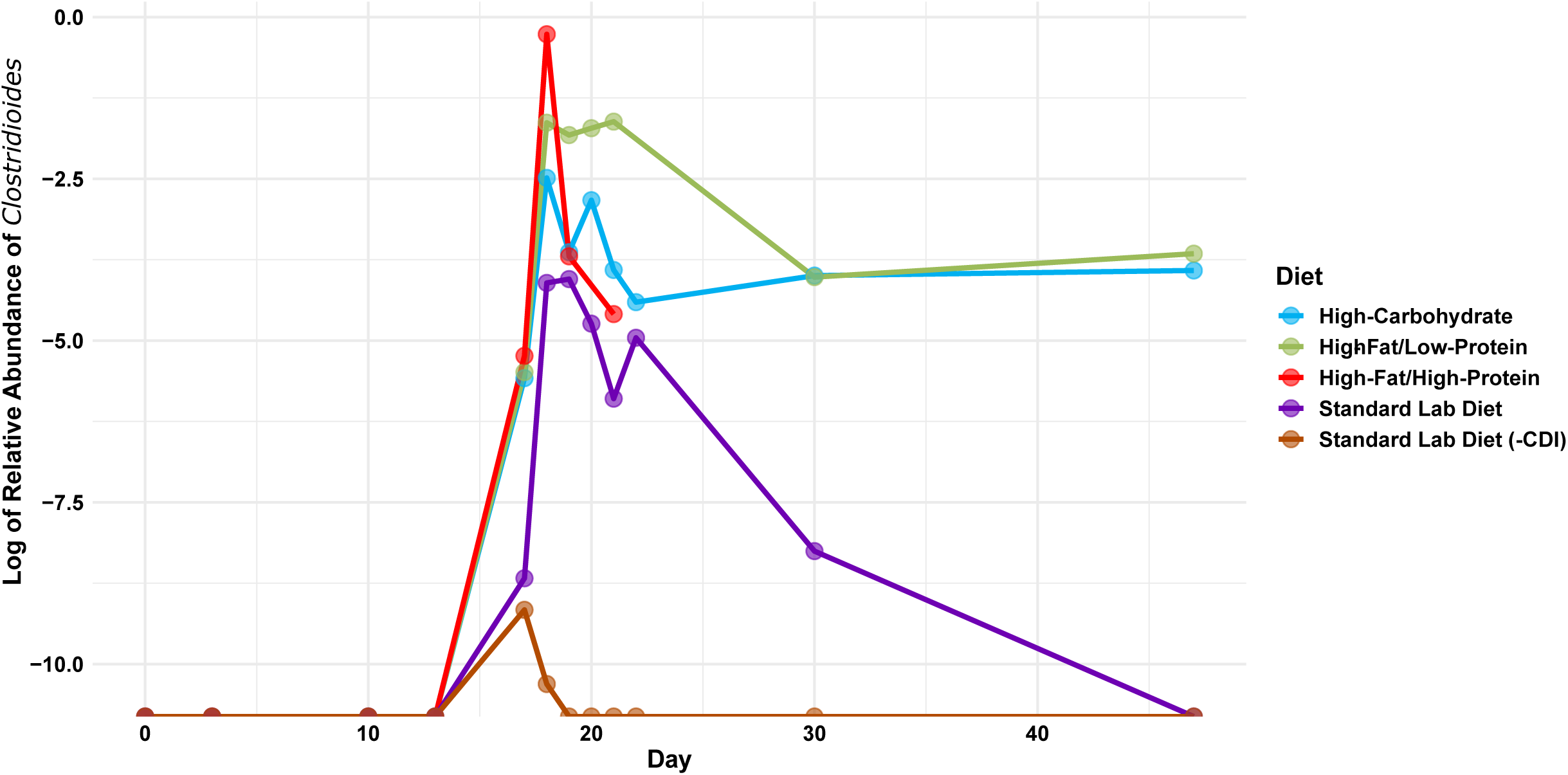
Log relative abundance of Clostridioides across experimental timeline. Log relative abundance of Clostridioides was calculated for uninfected mice fed a standard lab diet (orange), and infected mice fed a standard lab diet (purple, n = 5), high-carbohydrate diet (blue, n = 5), high-fat/low-protein diet (green, n = 5), or high-fat/high-protein diet (red, n = 5).

**File S1:**
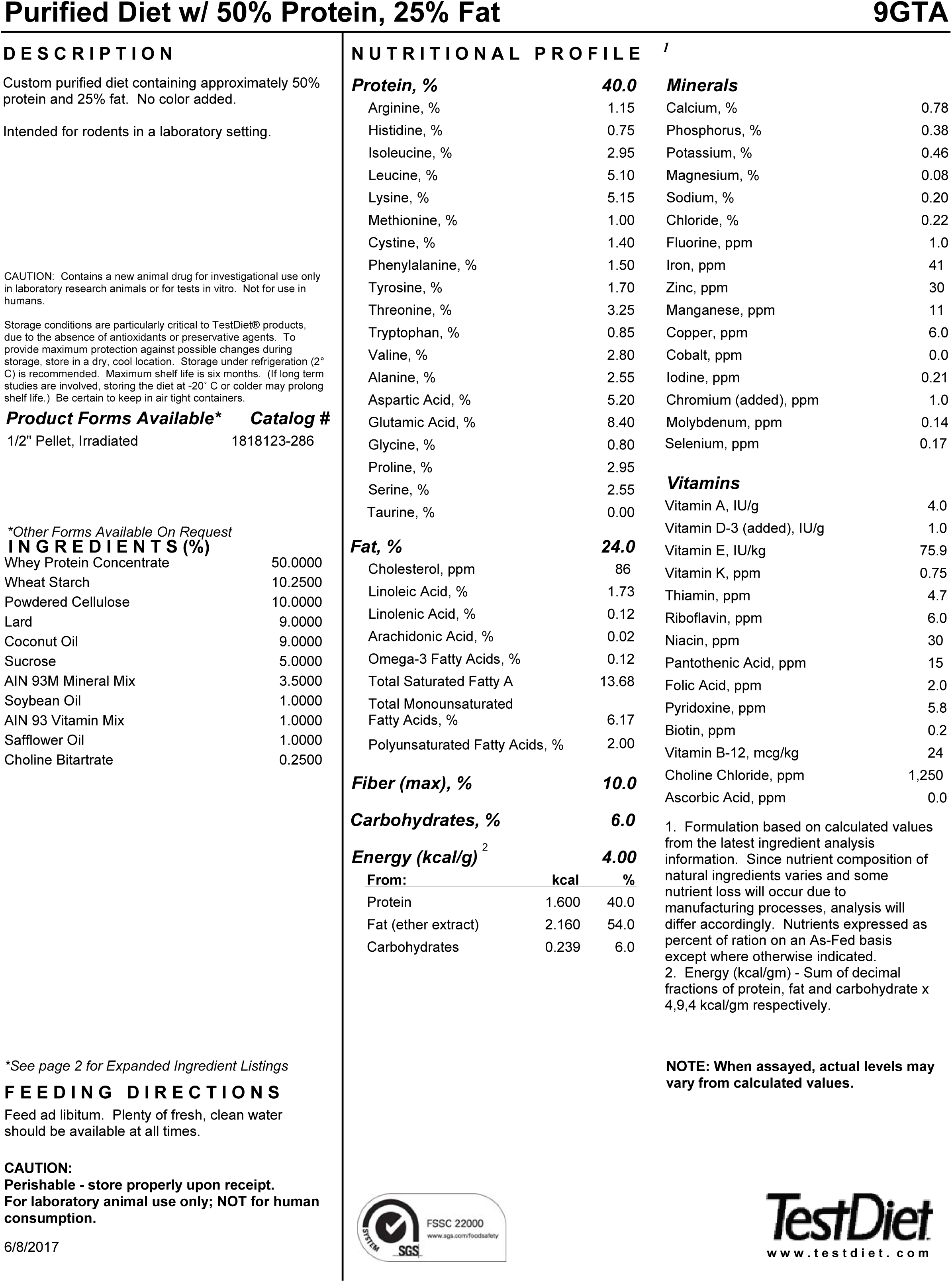
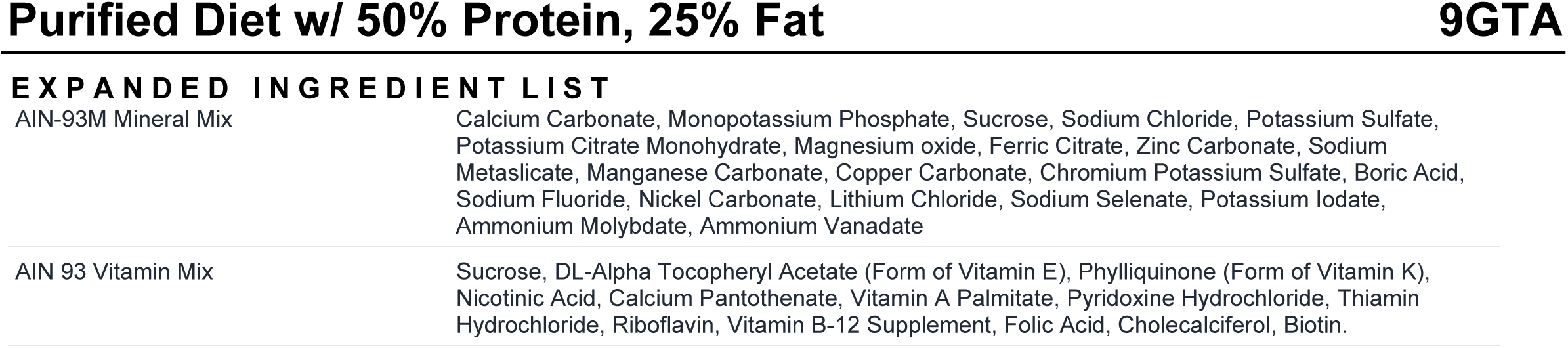
High-fat/high-protein diet. Freudenberg A, Petzke KJ, Klaus S. 2012. Comparison of high-protein diets and leucine supplementation in the prevention of metabolic syndrome and related disorders in mice. J Nutr Biochem 23:1524–1530.

**File S2:**
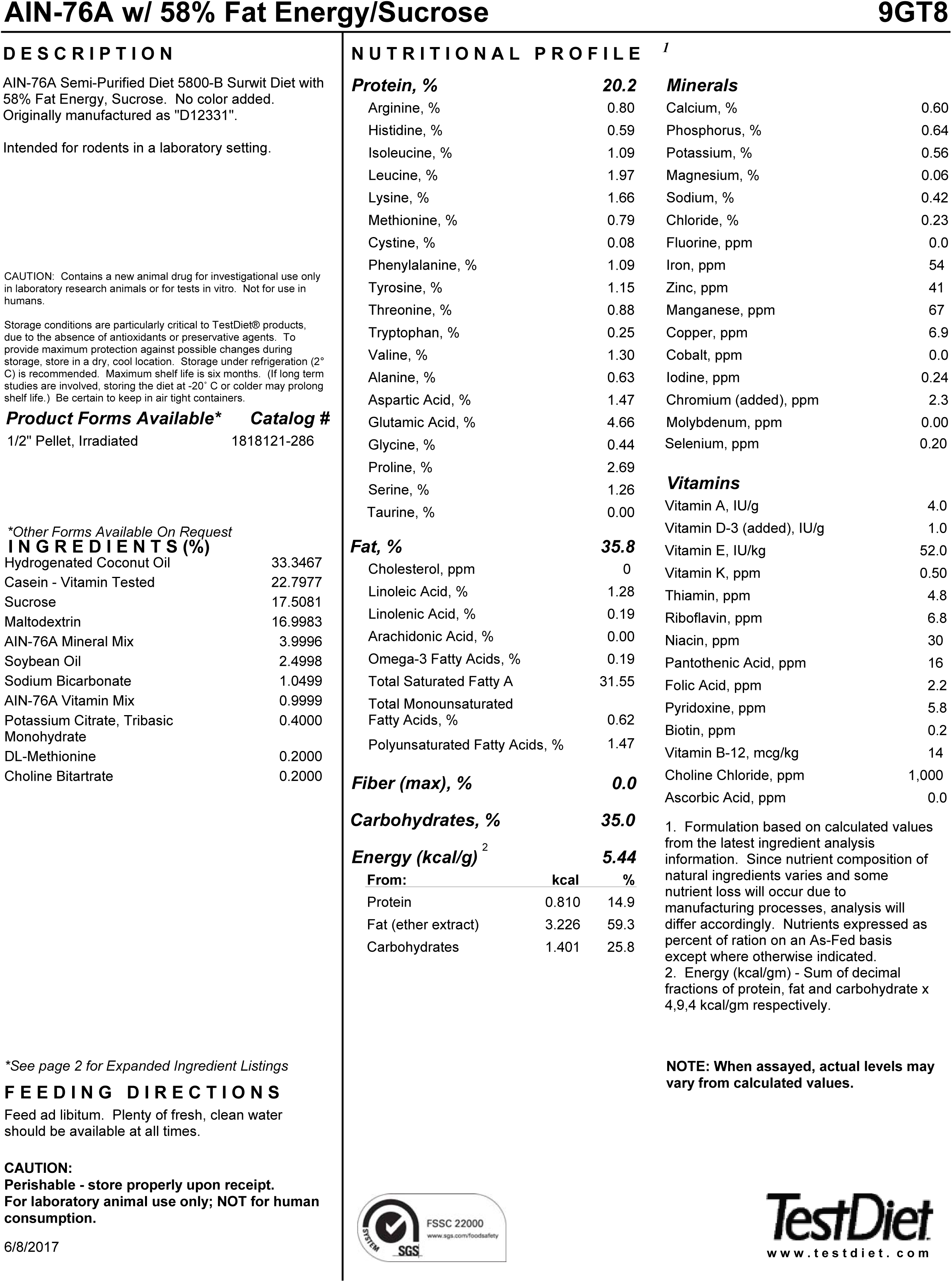
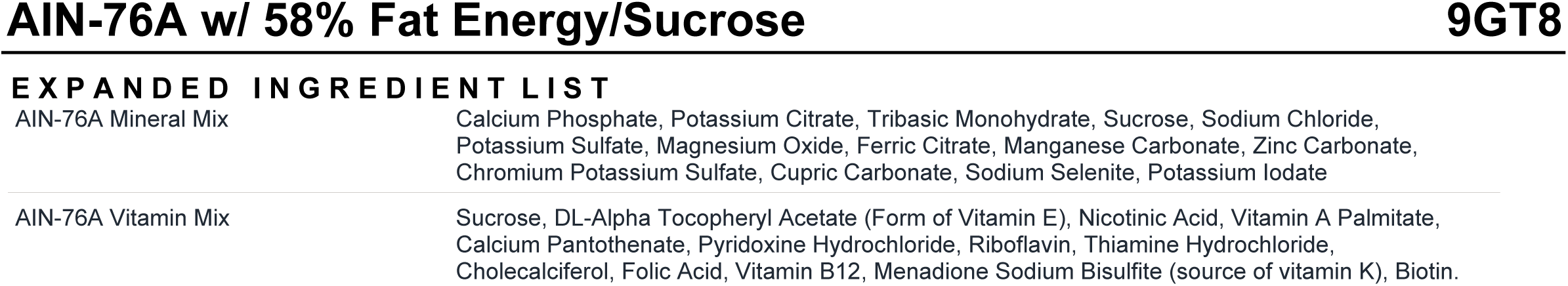
High-fat/low-protein diet.

**File S3:**
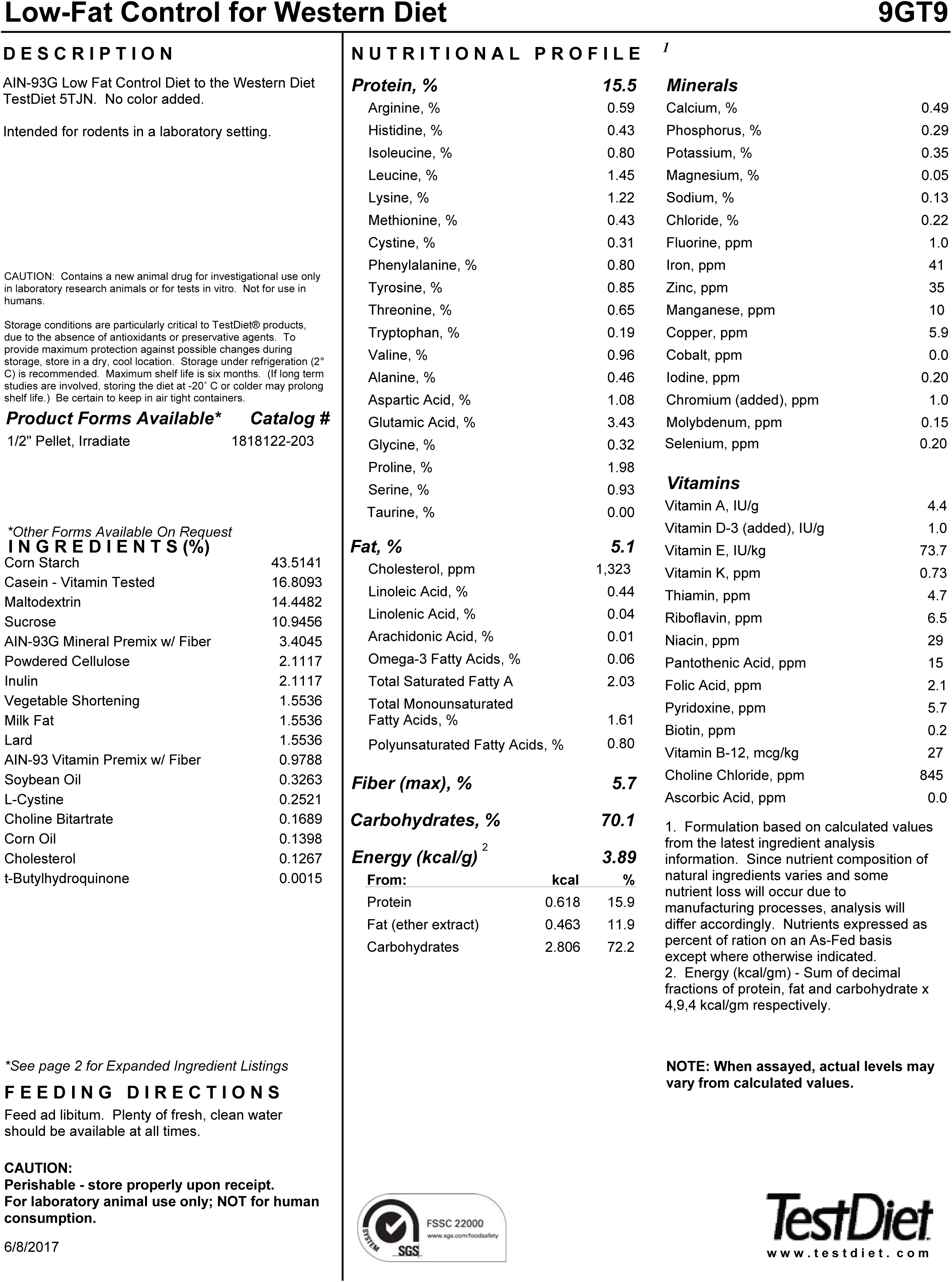
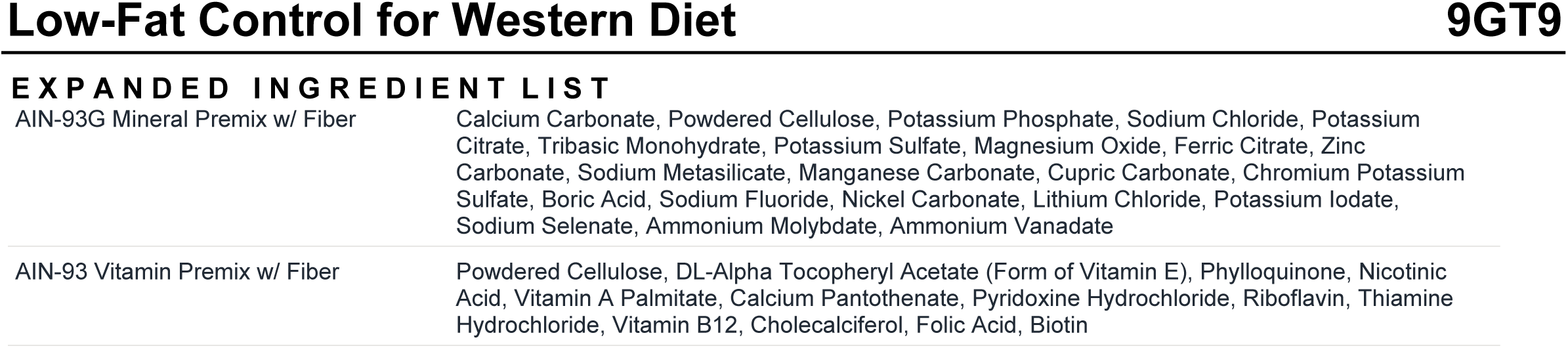
Low-fat/high-carbohydrate diet.

**Table S1:** Experimental setup for testing effect of diet on CDI mouse model.

| Diet | Purpose | Antibiotic<br>Cocktail | Clindamycin | *C. difficile* | % Kcal<br>Fat | % Kcal<br>Protein | % Kcal<br>Carbohydrate |
| --- | --- | --- | --- | --- | --- | --- | --- |
| Standard lab diet<br>+CDI<sup>1</sup> | - Control | + | + | + | 13.4 | 29.8 | 56.7 |
| Standard (-CDI)<sup>1</sup> | + Control | - | - | - | 13.4 | 29.8 | 56.7 |
| High-fat/high-protein diet<sup>2</sup> | Experimental | + | + | + | 54 | 40 | 6 |
| Low-fat/high-carbohydrate diet<sup>3</sup> | Experimental | + | + | + | 11.9 | 15.9 | 72.2 |
| High-fat/low-protein diet<sup>4</sup> | Experimental | + | + | + | 59.3 | 14.9 | 25.8 |

